# Flavonols modulate lateral root emergence by scavenging reactive oxygen species in *Arabidopsis thaliana*

**DOI:** 10.1101/2020.05.31.126557

**Authors:** Jordan M. Chapman, Gloria K. Muday

**Affiliations:** Biology department, Wake Forest University, Winston Salem, North Carolina, United States of America

**Keywords:** flavonoid, Arabidopsis thaliana, ROS, antioxidant, redox signaling, lateral root development

## Abstract

Flavonoids are plant-specific antioxidant compounds that modulate plant development, which include flavonols and anthocyanins subclasses. In *Arabidopsis thaliana*, mutants in genes encoding each step in the flavonoid biosynthetic pathway have been isolated. We used these mutants to examine the role of flavonols in initiation and emergence of lateral roots and asked whether this regulation occurs through scavenging ROS. The *tt4* mutants have a defect in the first committed step of flavonoid biosynthesis and have increased lateral root emergence. This phenotype was reversed by both genetic and chemical complementation. Using these flavonoid biosynthetic mutants, we eliminated roles for anthocyanins and the flavonols, quercetin and isorhamnetin, in controlling lateral root development. The *tt7*-2 mutant has a defect in a branchpoint enzyme blocking quercetin biosynthesis that led to elevated levels of kaempferol and reduced lateral roots. Kaempferol accumulated within lateral root primordia and was significantly increased in *tt7-2*. Thee data are consistent with kaempferol acting as a negative regulator of lateral root emergence. We examined ROS accumulation above and within the primordia using a general ROS sensor and identified increased signal above the primordia of the *tt4* and *tt7-2* mutants compared to wild type. Using a superoxide specific sensor, we detected a decrease in signal within the primordia of *tt7-2*, but not the *tt4* mutant, compared to wild type. Together, these results support a model in which increased level of kaempferol in *tt7*-2 leads to a reduction in superoxide concentration in the lateral root primordia thereby reducing ROS-stimulated lateral root emergence.

## Introduction

Flavonoids are a class of plant specialized metabolites with important functions in modulating development and stress responses (1, 2). There are multiple subclasses of flavonoids including chalcones, flavones, isoflavonoids, flavanones, flavonols, and anthocyanins (3). The pathway begins with the conversion of 4-Coumaroyl CoA and Malonyl CoA to naringenin chalcone. Further downstream, dihydroflavonols are produced and can be converted to either flavonols or anthocyanins. Flavonols and anthocyanins are two of the best studied flavonoid subclasses due to their ubiquitous presence in multiple species, their diverse functionality (4), and antioxidant capacity (5–8). *Arabidopsis thaliana* synthesizes three flavonols: kaempferol, quercetin, and isorhamnetin, which differ by the presence of a hydroxyl or methoxy group on their ring. The flavonoid biosynthesis pathway is well characterized in Arabidopsis with mutants isolated in the genes encoding each of the enzymes of the biosynthetic pathway (3).

Flavonoid biosynthetic mutants have been used to better understand the developmental role of flavonols and their role as signaling molecules. The Arabidopsis *transparent testa 4 (tt4)* mutant produces no flavonoids due to a mutation in the gene encoding the enzyme catalyzing the first committed step in the flavonoid biosynthesis pathway, chalcone synthase (CHS). Previous studies showed that *tt4* has increased root hair formation (9), ABA-induced stomatal closure (10), impaired gravity response (11), and greater sensitivity to environmental stress (12). Additional pathway mutants have been used to reveal which flavonols function in development, including identification of specific kaempferol derivatives that regulate leaf shape (13) and a role of quercetin in regulating gravitropic curvature and root hair initiation (9, 11). Using mutants impaired in flavonol biosynthesis in crop species, such as tomato, has also highlighted the role of flavonol metabolites in modulating environmentally-responsive signaling pathways, such as ABA-dependent guard cell signaling that induces stomatal closure (14) and temperature-impaired pollen viability and pollen tube growth (15). The following two mechanisms have been suggested to explain the controls of flavonols on plant development. Arabidopsis mutants with defects in flavonol synthesis have elevated levels of auxin transport, consistent with the absence of a negative regulator of auxin transport (11, 16, 17). Consistent with the antioxidant capabilities of flavonols, mutants that have impaired flavonol biosynthesis have higher levels of ROS in guard cells, root hairs, and pollen tubes (9, 10, 14, 15).

Flavonoids have been suggested to regulate root development, based on an increased number of lateral roots in the *tt4(2YY6)*(18) and *tt4-1* (19) mutants. The initiation and elongation of lateral roots is tightly regulated, with ROS being implicated in this process (20–22). Lateral roots initiate from founder cells in the pericycle that remained competent to divide during their transition from the root apical meristem to the differentiation zone of the root (23). After founder cells begin to divide asymmetrically, the lateral root primordium (LRP) develops through a series of eight stages (23, 24) where the LRP expands through the endodermis, cortex, and finally emerges through the epidermis. This process is regulated by mechanical signals (25–27), hormonal signals including auxin (25, 28–30), and biochemical signals such as ROS (20–22) or nitric oxide (31).

ROS can function as signaling molecules to regulate plant development, environmental responses, and hormone signaling (1, 32, 33). The low baseline levels of ROS allow local increases in ROS to act as important developmental signals. In Arabidopsis, ROS signaling has been implicated in stomatal closure (10, 14, 34, 35), pollen tube growth and development (15, 36–38), root hair elongation (9, 39), root gravitropism and phototropism (40–42), primary root elongation (43), the transition from cell proliferation to elongation in the root tip (44), and lateral root emergence (20–22). ROS signaling is also involved in the production of the casparian strip (45), and resistance to stress including hypoxia, salt stress, and ozone (46–48). Yet, ROS are also produced as a result of metabolism and stress (49, 50). Therefore, plant cells have elaborate mechanisms to keep ROS at low levels to prevent oxidative damage including the production of antioxidant enzymes (49, 51, 52) and specialized metabolites, such as flavonols (1).

This study examined the role of flavonols in modulating lateral root development and tested the hypothesis that flavonoids regulate this developmental process by scavenging ROS within the LRP. We quantified primordia and emerged lateral roots in a suite of flavonoid biosynthetic mutants, revealing that mutants which produce no flavonols had increased lateral root emergence. In contrast, a mutant that produces kaempferol at higher than wild type levels and produces no other flavonols had fewer emerged lateral roots than wild type, suggesting that kaempferol negatively regulates lateral root emergence. Consistent with an inhibitory effect of kaempferol, there is a negative correlation between the amount of kaempferol and the number of emerged lateral roots across multiple mutant lines. Using confocal microscopy and dyes that fluoresce upon binding flavonols or in response to oxidation by ROS, we find that in regions of roots with high levels of flavonols, there is decreased ROS abundance and in areas of high ROS, there are low concentrations of flavonols. The experiments in this manuscript support the model that flavonol-modulated root development is orchestrated by the scavenging of ROS within the lateral root primordia.

## Results

### Flavonoid mutants have predicted metabolite profiles

The flavonoid biosynthetic pathway is well characterized in Arabidopsis, with mutations mapped to the genes encoding each biosynthetic enzyme. The flavonoid biosynthetic pathway and the enzymes defective in these mutants are shown in Figure 1. We verified the metabolite profiles in the roots of the mutant alleles used in this study and under our specific growth conditions using liquid chromatography-mass spectroscopy (LC-MS) of aglycone flavonols using a standard curve and MS2 verification. Flavonol levels were quantified in Col-0 and six mutants with defects in genes encoding enzymes involved in flavonoid biosynthesis. The flavonol abundance relative to Col-0 for each mutant is shown in Figure 1 and the absolute value of each flavonol in nanomole per gram fresh weight (nmol/gfw) is reported in Table 1. In Col-0, naringenin and isorhamnetin were found at low concentrations, suggesting an efficient conversion of naringenin, an early pathway intermediate, into flavonols, and limited conversion of quercetin into isorhamnetin (Table 1).

**Table 1:** Quantification of flavonoids in roots of 7-day old seedlings by liquid chromatography mass spectrometry.

|  |  |  |  |  |  |
| --- | --- | --- | --- | --- | --- |
|  | Naringenin | Kaempferol | Quercetin | Isorhamnetin | Total |
| Col-0 (WT) | 0.39 ± 0.08 | 200.2 ± 22.06 | 354.6 ± 42.51 | 18.44 ± 6.06 | 573.7 ± 67.1 |
| *tt4-2* | 0.01 ± 0.003 | 16.3 ± 3.89** | 0.40 ± 0.09*** | N.D. | 16.6 ± 4.0*** |
| *tt4-11* | N.D. | N.D. | 0.01 ± 0.01*** | N.D. | 0.01 ± 0.01*** |
| *tt7-2* | 0.22 ± 0.02 | 363.3 ± 115.6*** | 2.60 ± 0.8*** | N.D. | 366.0 ± 128.8 |
| *fls1-3* | 0.58 ± 0.05 | 1.8 ± 0.8* | 5.03 ± 1.6 | 0.27 ± 0.10 | 7.8 ± 2.4*** |
| *omt-1* | 0.67 ± 0.10 | 276.9 ± 19.5 | 493.7 ± 55.52 | 6.3 ± 4.34 | 777.6 ± 73.2 |
| *tt3* | 0.19 ± 0.01 | 162.3 ± 36.19 | 274.49 ± 63.69 | 0.01 ± 0.003 | 436.9 ± 99.4 |
| Statistics were measured by a 2-way ANOVA with a Tukey post-hoc test over three separate replicates with an n= 4-13 (*p<0.05, ** p<0.005, *** p<0.0001). N.D indicates values that were not detected because they were below the lowest concentration on the standard curve. |  |  |  |  |  |

**Figure 1:**
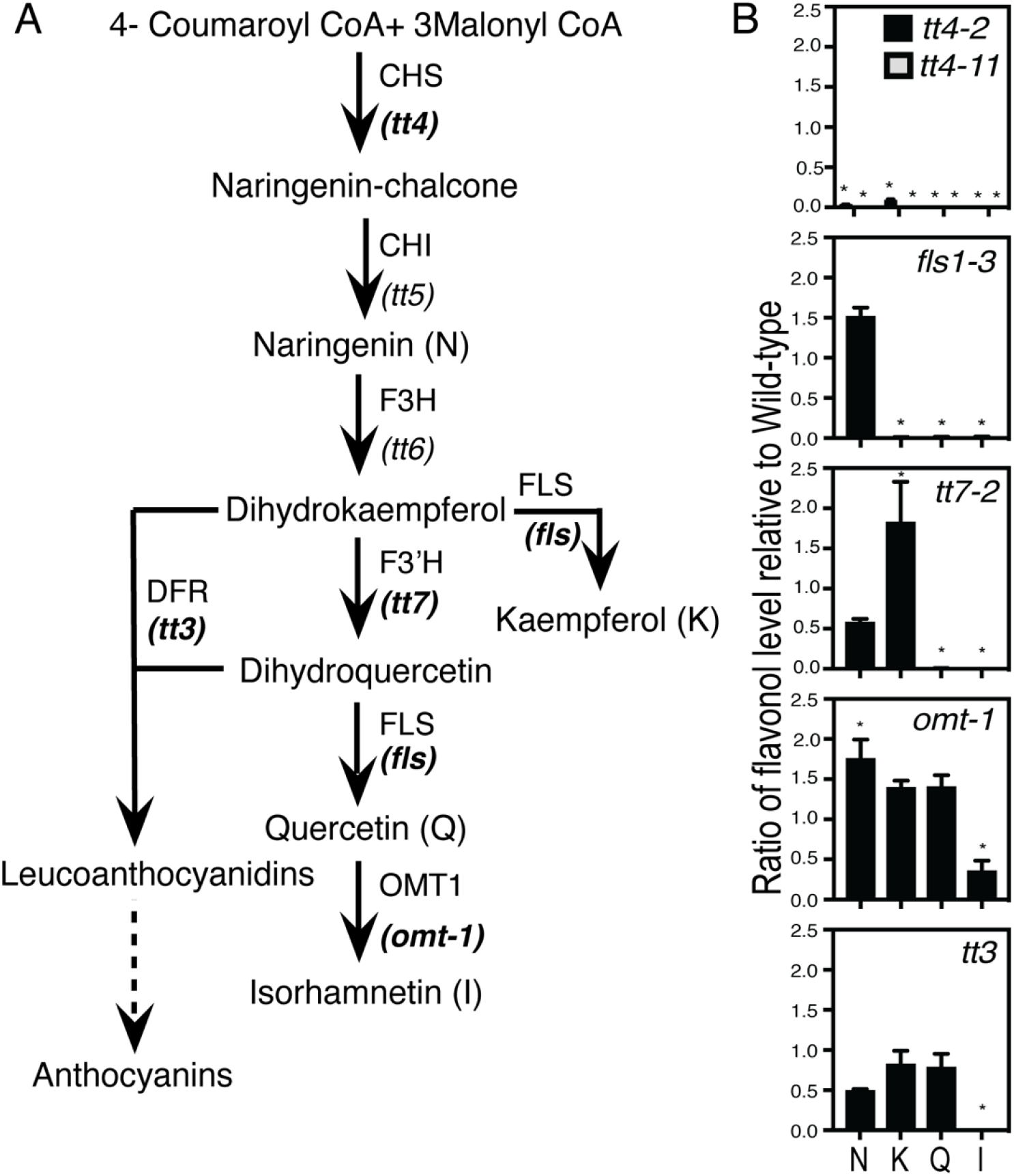
Flavonoid biosynthesis mutants have expected flavonol profiles. (A)The flavonoid biosynthesis pathway in Arabidopsis has mutants at each step indicated in italics under the enzyme. Mutants in bold were used in the experiments in this manuscript. (B) Flavonol levels were determined by LC-MS and are reported in nmole/gfw (gram fresh weight) and a ratio of the mutant wild type levels were calculated and reported above. Statistics were determined using a two-way ANOVA followed by a Tukey post-hoc test. * Indicates a p< 0.05. Data was collected from three separate experiments with total n of Col-0 (n=22), *fls1-3* (n=4), *tt4-2* (n=12), *tt4-11* (n=6), *tt7-2* (n=5), *omt-1* (n=7), and *tt3* (n=11).

The flavonol accumulation profiles in the mutants are consistent with mutant enzyme function in the biochemical pathway (Figure 1). The *tt4-2* and *tt4-11* mutants have defects in the gene encoding CHS, while the *fls1-3* mutant has a defect in the gene encoding flavonol synthase1 (FLS1), which is the predominant FLS isozyme catalyzing the final step of flavonol biosynthesis (53, 54). The *tt4-2, tt4-11* and *fls1-3* mutants have background levels of the 3 flavonols: kaempferol, quercetin, and isorhamnetin. The *fls1-3* mutant has significantly increased levels of naringenin, due to its inability to produce downstream products. The *tt7-2* mutant has a defect in the gene encoding the branch point enzyme Flavonoid 3’-Hydroxylase (F3’H). It produces no quercetin but has 1.8-fold higher levels of kaempferol. The *omt-1* mutant, which cannot convert quercetin to isorhamnetin, produces significantly reduced levels of isorhamnetin. However, the levels of kaempferol and quercetin in *omt-1* are not significantly different from Col-0 likely due to low flux towards isorhamnetin in roots. The *tt3* mutant has a defect in the gene encoding dihydroflavonol 4-reductase (DFR), which converts dihydroxyflavonols to anthocyanins. It produces wild type levels of flavonols in roots. This is likely due to the absence of DFR transcript and protein in roots of Col-0 (11, 55). Our flavonol metabolite profiling reveals that the specific alleles of these biosynthetic mutants have flavonoid profiles in roots consistent with their genetic defect.

### Mutants with impaired flavonol synthesis have increased lateral root number

To evaluate root development in the absence of all flavonols, lateral root number of 8-day old seedlings was quantified in Col-0 and three mutants that do not produce flavonols. Representative images of the lateral root phenotypes of *tt4-2, tt4-11*, and *fls1-3* are shown in Figure 2A. There are significantly more lateral roots in both *tt4* alleles as compared to the wild type, consistent with a prior report using the *tt4(2YY6)* allele (18). The *fls1-3* mutant synthesizes all the pathway intermediates but makes no flavonols and also has significantly more lateral roots than Col-0. This result suggests that the flavonols, rather than their precursors, limit the formation and/or emergence of lateral roots in these flavonol biosynthesis mutants.

**Figure 2:**
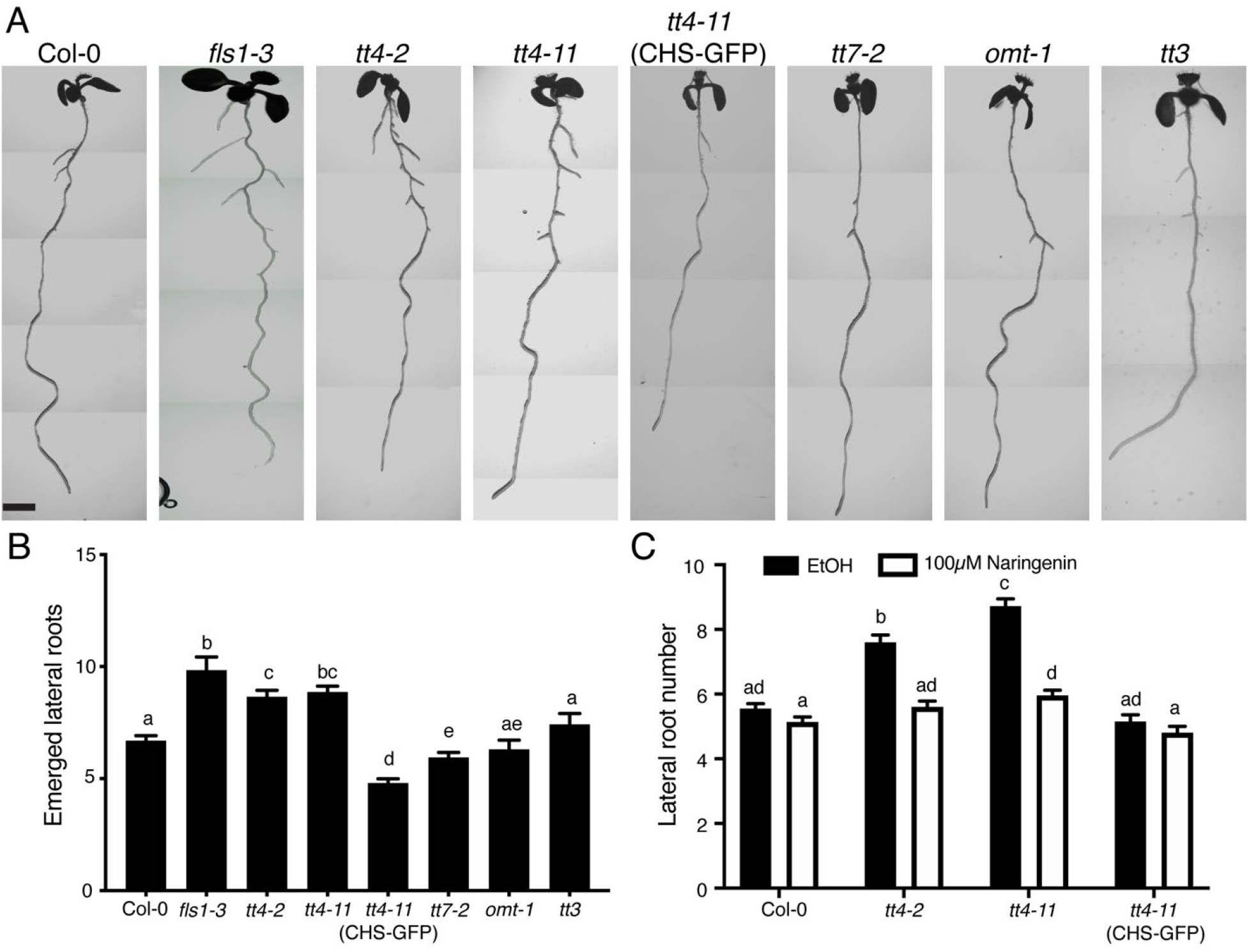
Mutants in the flavonoid biosynthetic pathway reveal that kaempferol is a negative regulator of lateral root emergence. (A) Representative images of each mutant were taken using a Zeiss Axiozoom microscope. Scale bar: 2mm. (B) Lateral root number was quantified over four replicates with total number of seedlings: Col-0 (n=161), *fls1-3* (n=40), *tt4-2* (n=122), *tt4-11* (n=67), CHS-GFP (n=69), *tt7-2* (n=121), *omt-1* (n=45), and *tt3* (n=58). Statistics were performed using a one-way ANOVA with p<0.05. (C) Lateral root number was evaluated after a three-day treatment with 100 μM naringenin. The average and SE are from four independent experiments with the following total n: Col-0 (EtOH: n= 89; Naringenin: n=89), *tt4-2* (EtOH: n= 50; Naringenin: n=50), *tt4-11* (EtOH: n= 59; Naringenin: n=64), and CHS-GFP (EtOH: n= 45; Naringenin: n=45). Statistics were measured using a two-way ANOVA followed by a Tukey post-hoc test p< 0.05. Bars with the same letter represent no statistical difference, while different letters indicate values that are significantly different.

### Genetic and chemical complementation of *tt4-2* and *tt4-11* reduce lateral root number to wild type levels

The *tt4-11* mutant was complemented both genetically and chemically to verify the loss of function of *CHS* resulted in the increased lateral root phenotype. The *tt4-11* mutant was genetically complemented with the *CHSpro::CHS-GFP* transgene: *tt4-11*(CHS-GFP) and lateral root number was quantified. The genetic complementation of *tt4-11* with CHS-GFP resulted in significantly fewer lateral roots compared to the untransformed *tt4-11* mutant and Col-0 (Fig 2B). We also chemically complemented *tt4-2* and *tt4-11* with the flavonol precursor naringenin, a pathway intermediate produced downstream of CHS. 5-day old seedlings of Col-0, the two *tt4* alleles, and *tt4-11* (CHS-GFP) were transferred to media supplemented with 100 μM naringenin for three days before the number of lateral roots was quantified. The lateral root numbers in *tt4-2* and *tt4-11* were significantly reduced by naringenin treatment to wild type levels. In Col-0 and the *tt4-11*(CHS-GFP) line, which endogenously produce naringenin, the number of lateral roots was not significantly different in the presence or absence of added naringenin (Figure 2C). The absence of an effect with naringenin supplementation in Col-0 is consistent with the conversion of naringenin to dihydrokaempferol being rate limiting. These complementation results are consistent with impaired flavonol synthesis leading to an increased number of lateral roots.

### Flavonoid biosynthesis mutants reveal the pathway intermediates that regulate root development

Lateral root development was evaluated in 8-day old seedlings in mutants at multiple steps in the flavonoid biosynthetic pathway to determine whether a specific flavonol(s) functions to modulate this development. Representative images of these roots are shown in Figure 2A and the number of emerged lateral roots was quantified and are reported in Figure 2B. The primary root length for each mutant was also measured and there were no significant differences in length in *tt4-2, tt4-11, fls1-3, tt7-2*, and *tt3* mutants compared to the wild type (Figure S1), therefore the effect on lateral root development was independent of root length.

The lateral root number of the *omt-1* and *tt3* mutants were not significantly different from wild type, consistent with the absence of root developmental role of isorhamnetin and other downstream molecules in the flavonoid pathway, including anthocyanins. The mutant *tt7-2*, which has higher levels of kaempferol and for which quercetin and isorhamnetin are not synthesized, has a decreased number of lateral roots (Figure 2B). The root developmental pattern of *tt7-2* suggests either kaempferol specifically inhibits lateral root formation/emergence or that local concentration changes of kaempferol regulate development.

### Flavonol mutants show altered lateral root emergence and initiation

We asked whether the increase in lateral root number in the *tt4* alleles and decrease in *tt7-2* were due to modulation of lateral root emergence, lateral root initiation, or a combination of both. Lateral root initiation begins when founder cells exhibit an asymmetric division (stage 1), which is then followed by a precise series of divisions to form LRP that ultimately move through the epidermis as emerged lateral roots (stage 8) (23). To determine the LR developmental stage at which lateral roots were impaired in *tt7-2*, we quantified the number of LRP between stage 3 and stage 8 and emerged lateral roots in *tt4-2, fls1-3*, and *tt7-2* (Figure 3A) and in *tt4-11, omt-1*, and CHS-GFP (*tt4-11*) (Figure S2). The number of LRP in *tt4-2, tt4-11*, and *fls1-3* mutants were not significantly different from Col-0. In contrast, these genotypes all had a significantly higher number of emerged lateral roots than Col-0. The *tt7-2* mutant had slightly increased LRP compared to Col-0, which as significant by a Student’s t-test, but not by a more stringent 2-way ANOVA as reported in Figure 3A. In contrast, the *tt7-2* mutant has a significantly decreased number of emerged lateral roots relative to Col-0 by all statistical tests (Figure 3A). This result indicates that in *tt4-2, tt4-11* and *fls1-3* mutants root emergence is enhanced, while in *tt7-2* emergence is impaired.

**Figure 3:**
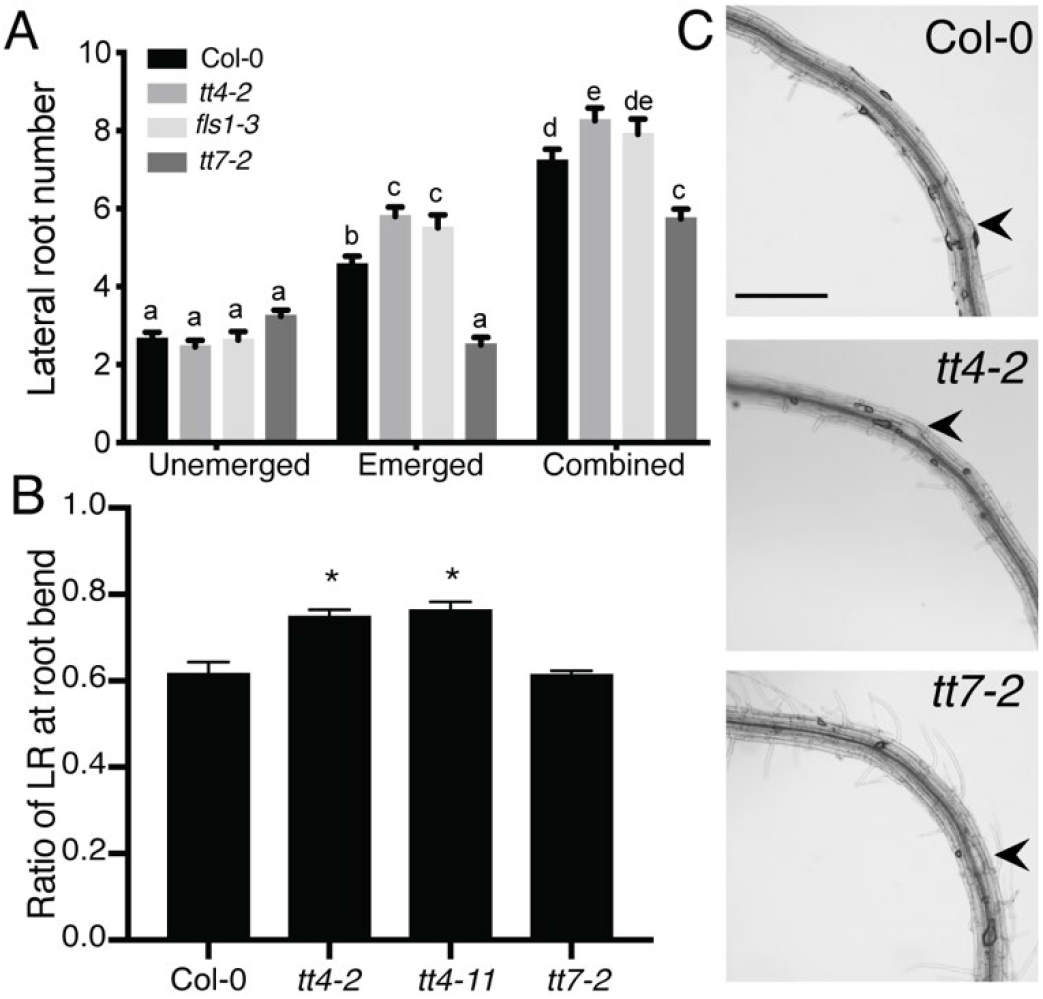
Flavonol mutants have altered lateral root emergence. (A) The number of lateral primordia (Stages 4-7) and emerged lateral roots, and the combined totals were quantified in cleared 7-day old seedlings. The average and SE from four separate experiments with a total n=60-68. Statistical differences were determined with a one-way ANOVA with p<0.05. Bars with the same letter indicate no statistical significance while bars with different letters are statistically significant from one another. (B) Roots were bent after five days of growth to a 90°-angle and determination of lateral root formation was done two days later. The average and SE from five separate experiments with a total n: (Col-0 n=71; *tt4-2* n=44; *tt4-11* n= 72; *tt7-2* n=70) is reported. Statistical differences were determined using a one-way ANOVA with p<0.0001. (C) Representative images of lateral root primordia in bent roots indicated by black arrows. Scale bar = 50μm.

In Arabidopsis, a 90-degree bend in the primary root leads to the production of a lateral root at the position of the bend (26, 27, 56). We took 5-day old seedlings and reoriented them with a 90-degree angle at a 2mm distance from the root tip. The formation of a LRP or emerged LR was evaluated after 48 hours with representative images shown in Figure 3C. The number of roots with lateral roots (emerged or unemerged) at the bend position was divided by the total number of roots to derive the ratio reported in Figure 3B. The ratios calculated in mutants were compared to wild type. Col-0 and *tt7-2* formed a lateral root in 61% of roots that were bent. The two *tt4* mutant alleles, *tt4-2* and *tt4-11*, formed significantly more lateral roots than wild type, with 75 and 76% of the bent roots initiating lateral roots, respectively. The increase in lateral root initiation in the *tt4* mutants likely contributes to the increased lateral root phenotype. These data indicate that the decrease in lateral root number in *tt7-2* is likely due to decreased emergence, not a decrease in lateral root initiation.

### Total flavonol level negatively correlates with lateral root number

To understand why *tt7-2* has an opposite lateral root phenotype compared to the flavonol-deficient mutants, we asked whether the number of emerged lateral roots was predicted by the amount of either quercetin, kaempferol, or the total flavonol level. We plotted the average number of emerged lateral roots in wild type and mutants as a function of either the quercetin or kaempferol concentration (Figure 4A), or total flavonol levels (Figure S3), followed by a linear regression analysis. While there was no significant correlation between the number of emerged lateral roots and the levels of quercetin, there was a significant negative correlation between the number of lateral roots and the level of both kaempferol and total flavonols. These data suggest the level of kaempferol and the total flavonol concentration is pertinent to lateral root emergence, leading us to ask whether the site of accumulation of kaempferol or just the amount of this flavonol controls lateral root formation.

**Figure 4:**
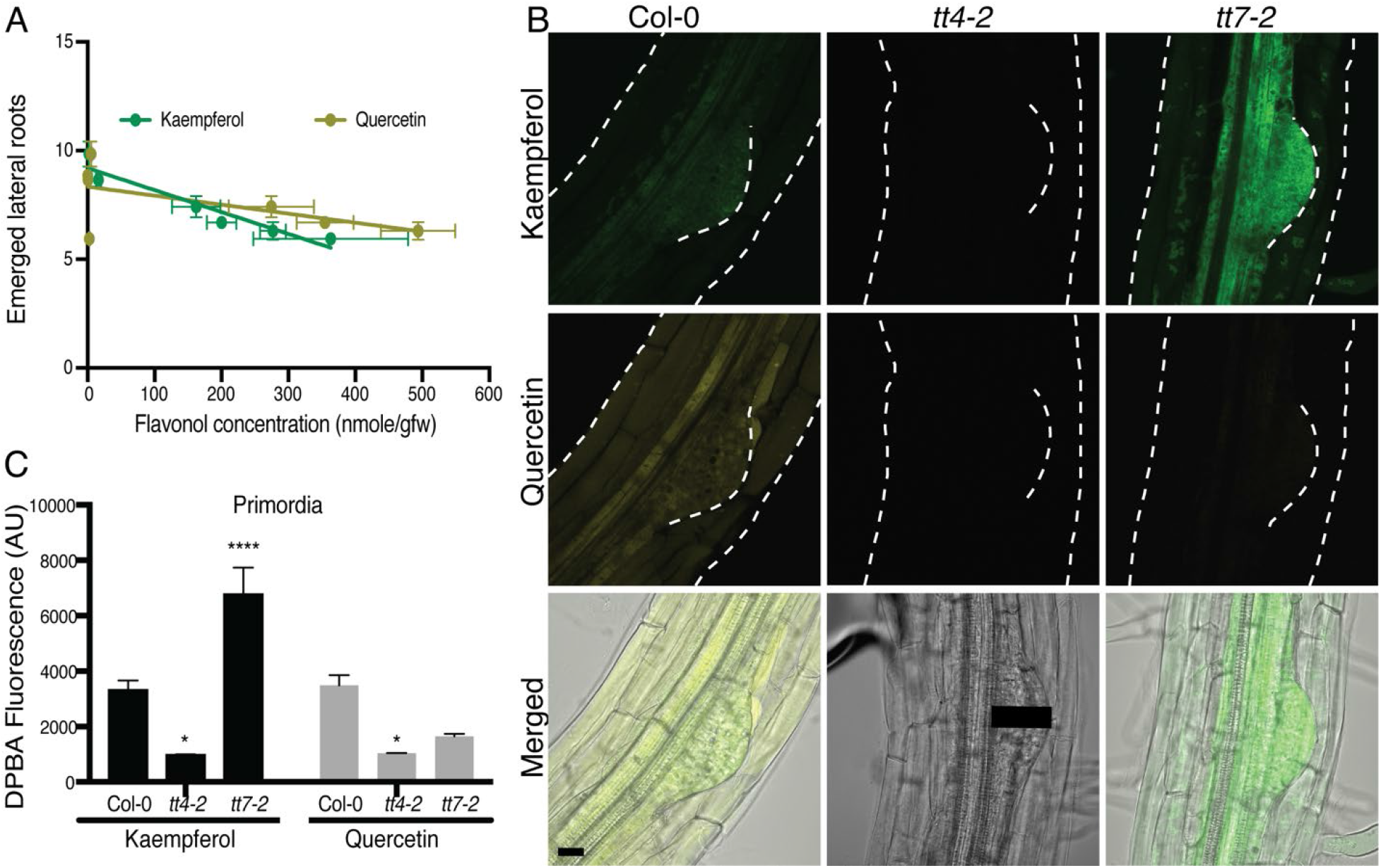
Kaempferol accumulates in lateral root primordia. (A) The number of emerged lateral roots were plotted as a function of concentration of kaempferol or quercetin for wild type and the suite of flavonol mutants. Results from a linear regression are drawn and the p value and R square are: kaempferol (p=0.0032, R^2^=0.9094) and Quercetin (p=0.1644, R^2^=0.3465). Kaempferol has a significantly negative correlation with emerged lateral root number. (B) DPBA fluorescence intensity was quantified for K-DPBA and Q-DPBA using a 100 pixel line (11.5μm wide) drawn from the vasculature to the tip of the primordia, as represented by the black line on the *tt4-2* DIC image. The average fluorescence of multiple roots taken at identical gain, laser intensity, and pinhole settings are shown. Statistics were calculated based on a two-way ANOVA with a Tukey post-hoc test with a total n: Col-0 (n=23), *tt4-2* (n=13), *tt7-2* (n=22). Mutant values were compared to wild type (* p<0.05, **** p<0.0001). (C) Representative images of 8-day old seedlings stained with DPBA are shown with K-DPBA (green) and Q-DPBA (yellow) imaged using two channel mode. Scale bar: 50μm.

### Flavonols accumulate within and around developing lateral root primordia

To understand how flavonols regulate lateral root development, we asked where kaempferol and quercetin accumulate in LRP and emerging lateral roots. We stained roots of 8-day old Arabidopsis seedlings with diphenylboric acid 2-aminoethyl ester (DPBA), a dye which becomes fluorescent upon binding kaempferol and quercetin (11, 57). The emission spectra resulting from DBPA-bound kaempferol (K-DPBA) can be resolved from DPBA-bound quercetin (Q-DPBA) and these two fluorescent signals can be quantified by laser scanning confocal microscopy (LSCM) (11).

Confocal images of DPBA stained Col-0 with K-DPBA shown in green and Q-DPBA shown in yellow, illustrate the accumulation of these two compounds in LRP and surrounding tissues (Figure 4B). To verify the specificity of DPBA for recognition of flavonols in mature regions of the root, several mutants were used. The *tt4-2* and *tt4-11* mutants, which produce neither flavonol, have only background levels of K- and Q-DPBA fluorescence (Figure 4B). In the *tt7-2* mutant, K-DPBA fluorescence is evident, with Q-DPBA signal at background levels consistent with the flavonol profile in Figure 1. In Col-0, both K- and Q-DPBA fluorescent signals are found in pericycle cells that give rise to LRP and within LRP (Figure 4B). In *tt7-2*, there is a striking apparent increase in levels of K-DPBA in primordia.

To determine whether the K-DPBA signal is consistently higher within LRP of *tt7-2* across multiple samples, the average fluorescence was quantified using a line drawn from the vasculature to the edge of the primordium through the middle of the LRP, as illustrated with the black line in Figure 4B. The vasculature is evident in differential interference contrast (DIC) images as the tissue directly below the pericycle. A line width of 11.5μM was used to account for any variance between flavonol accumulation that may occur across the width of the LRP. The K-DPBA and Q-DPBA fluorescence are found within primordia of Col-0, and both signals are at background levels in *tt4-2*. In *tt7-2*, the K-DPBA signal in lateral root primordia is significantly higher than Col-0 (p < 0.0001), while Q-DPBA fluorescence is at background levels and is not statistically different from those of *tt4-2*, consistent with the absence of quercetin (Figure 4B). The 2-fold increase in K-DPBA signal in primordia is similar to the 2-fold increase of kaempferol concentration detected by LC-MS (Figure 1). This pattern suggests that kaempferol accumulation in lateral root primordia is appropriately positioned to modulate lateral root development.

### Flavonol biosynthetic enzymes reporter fusions are expressed in pericycle cells and emerged lateral roots

Flavonols accumulated in founder cells and LRP, leading us to ask where their biosynthetic machinery localizes. We used two fluorescent protein constructs, *CHSpro:CHS:GFP* (CHS-GFP), and *FLS1pro:FLS1:GFP* (FLS-GFP) (11, 13) to visualize by LSCM where these key enzymes accumulate during lateral root development. Maximum intensity projections of representative images of LRP and emerged lateral roots expressing CHS-GFP and FLS-GFP are shown in Figure 5. GFP fluorescence of both reporters was detected in the pericycle cell files that give rise to lateral roots and at the base and the apical tip of emerged lateral roots (Figure 5). CHS-GFP signal was not detected in the LRP, while FLS-GFP signal at the tip of the LRP parallels the signal at the tip of emerged lateral roots. These data are consistent with flavonol synthesis in pericycle cells prior to formation of LRP or with a precursor for flavonols being transported to primordia for the production of flavonols. Prior reports indicates that naringenin can move long distances in roots, consistent with this later possibility (58, 59).

**Figure 5:**
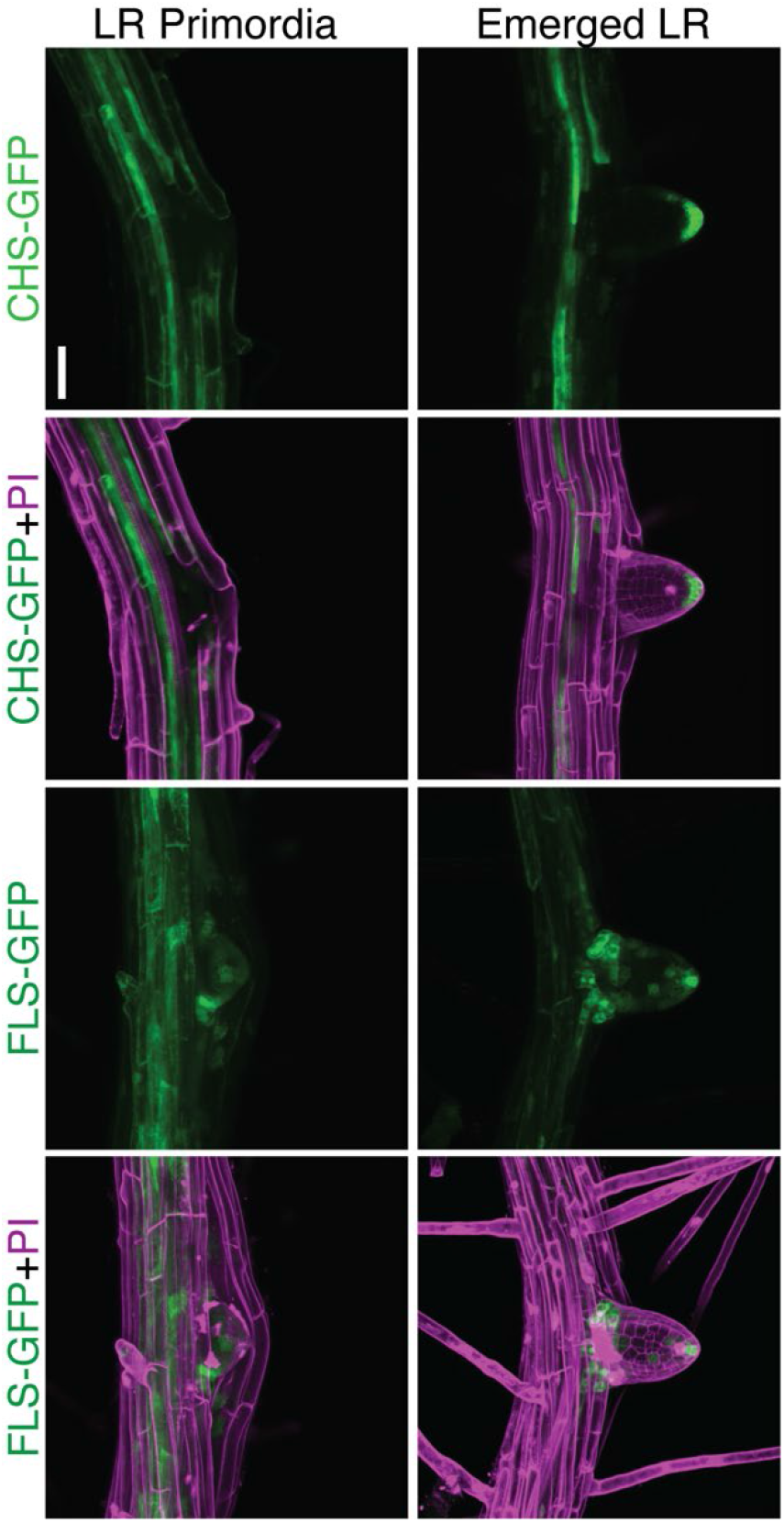
*CHSpro:CHS-GFP* and *FLSpro:FLS-GFP* accumulate in the emerged lateral root. CHS-GFP and FLS-GFP 8-day old seedlings were stained with propidium iodide shown in magenta. Representative images are from three separate experiments with identical gain, laser intensity, and pinhole. Scale bar: 50μm.

### The absence of flavonols leads to increased levels of ROS that can be rescued with ascorbic acid

We asked if the mechanism by which flavonol-deficient mutants have elevated lateral roots is through increases in ROS signaling, due to the absence of flavonol antioxidants. To test this hypothesis we used ascorbic acid, a ROS scavenger, to ask if the enhanced lateral root phenotype in the *tt4* mutant was reversed by antioxidant treatment. Seedlings were grown on MS media for 5 days and then transferred to media containing 500 μM ascorbic acid for 3 days before lateral root number was quantified. Treatment with ascorbic acid significantly reduced the lateral root number in both *tt4* mutant alleles to levels similar to or less than wild type, consistent with elevated ROS driving the increased lateral root emergence (Figure 6A). In contrast, the ascorbic acid treatment had no significant effect on Col-0 and *tt7-2*, which suggests that synthesis of a kaempferol antioxidant is sufficient to eliminate the effect of exogenous antioxidant treatment.

**Figure 6:**
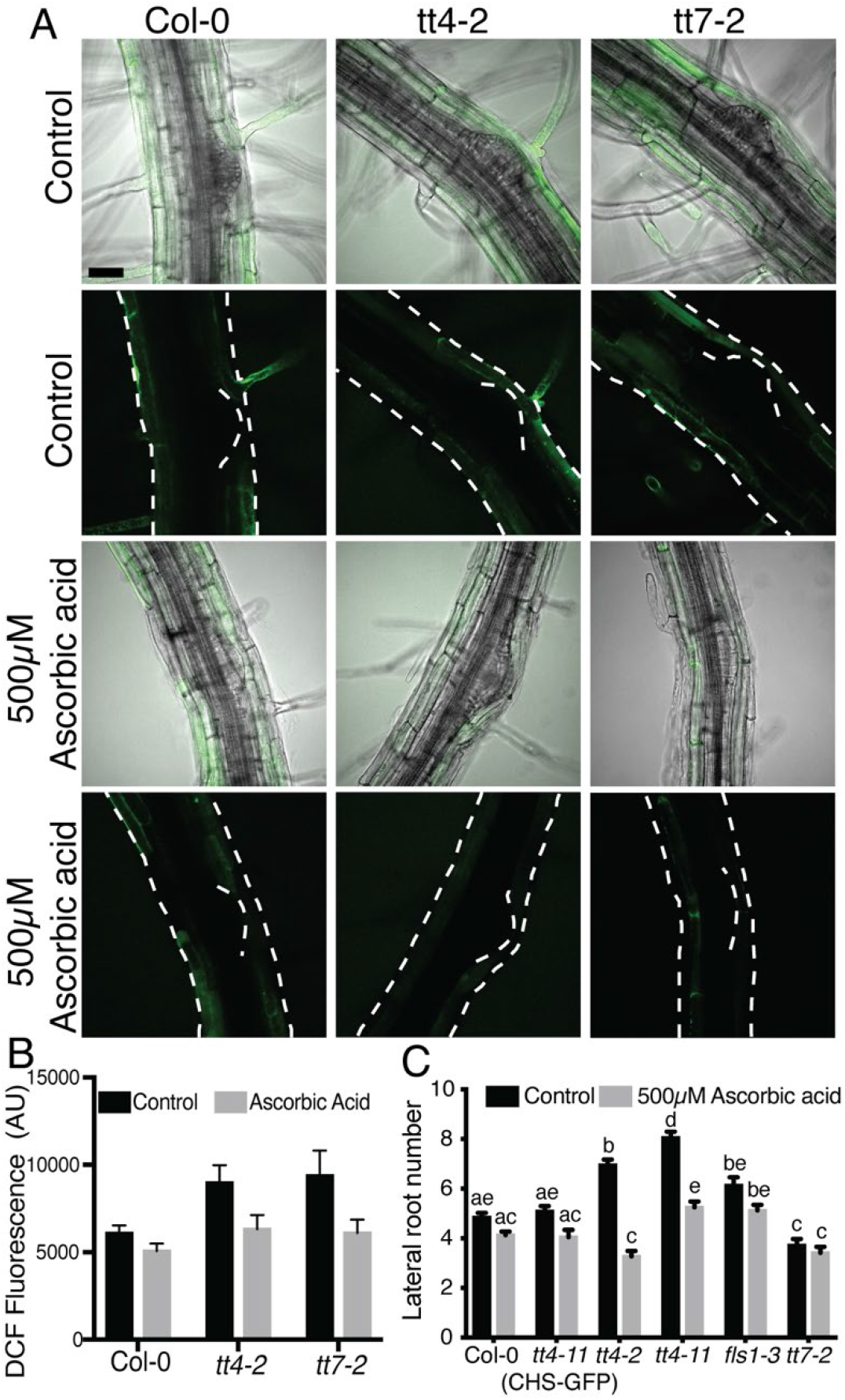
Treatment with ascorbic acid reduces lateral root number and DCF signal in *tt4-2*. (A) Lateral root number was evaluated after a 3-day treatment with 500 μM ascorbic acid. Data summarized from four independent experiments with varying total numbers: Col-0 (control: n= 93; ascorbic acid: n=89), *tt4-11*(CHS-GFP) (control: n= 43; ascorbic acid: n=36), *tt4-2* (control: n= 74; ascorbic acid: n=39), *tt4-11* (control: n= 48; ascorbic acid: n=55), *fls1-2* (control: n= 44; ascorbic acid: n=40) and *tt7-2* (control: n= 44; ascorbic acid: n=38). (B) 8-day old seedlings were stained with DCF and immediately imaged at the same laser power, gain, and pinhole for three separate experiments. Lateral root primordia were stage 4 and older due to limited stain uptake in primordia younger than stage 4. Data is summarized with a total n: Col-0 (control: n=13; ascorbic acid: n=19), *tt4-2* (control: n=15; ascorbic acid n=21), *tt7-2* (control: n=18; ascorbic acid: n=17). Statistics were measured using a two-way ANOVA with a post-hoc Tukey test. Bars with the same letter represent no statistical difference, while different letters indicate statistical significance with a p< 0.05.

To highlight where ROS accumulate across developing lateral roots and to examine the effect of ascorbic acid treatment on ROS levels, we examined levels of ROS in flavonol-deficient mutants in the absence and presence of ascorbic acid. We used a general ROS sensor, 2’,7’-dichlorodihydro-fluorescein diacetate (CM H_2_DCF-DA) and LSCM to visualize ROS in and around LRP. CM H_2_DCF-DA enters the cell through the plasma membrane, where the diacetate group is cleaved by intracellular esterases. The released DCF will fluoresce upon oxidation by ROS. DCF has a transient signal, which leads to fluorescence increases upon oxidation, but it can then be reduced and lose its fluorescence (60). DCF fluorescence accumulated in epidermal and cortical tissues above the lateral root primordia in wild type. We quantified DCF signal in untreated roots using a region of interest over the LRP. The *tt4-2* and *tt7-2* mutants both had significantly higher DCF signal above LRP than wild type (Figure 6B-C). Ascorbic acid treatment of Col-0 led to a reduction of DCF signal over the lateral root primordia. Similarly, treatment of *tt4-2* and *tt7-2* with ascorbic acid reduced DCF signal in this region to levels not significantly different from untreated Col-0, consistent with ascorbic acid acting as an antioxidant. Interestingly, DCF signal was not observed inside the lateral root primordia, where we see high levels of flavonols.

To determine whether the absence of DCF signal in the primordia is due to the inability of dye to be taken up in the primordia, we used the dye fluorescein diacetate (FDA) as a control, because it has a similar structure to DCF, but is not redox sensitive. LRP of stage 4 and later were able to take up FDA (Supplemental Figure 4) even though they showed no DCF signal. Lateral root primordia of stage 1-3 had neither FDA nor DCF signal, presumably because dye uptake is limited at these earlier developmental stages. The low level of DCF signal in primordia of wild type and flavonol mutants, even though FDA signal was detected, suggests ROS is present at low levels in the LRP. Although DCF is likely taken up in primordia, the transient nature of the signal diminishes our ability to evaluate ROS levels within the LRP.

### The *tt7-2* mutant has deceased superoxide (DHE) signal in the primordia compared to wild type

We used the sensor dihydroethidium, DHE, to monitor the accumulation of superoxide in wild type, *tt4-2*, and *tt7-2*. In all three genotypes, the DHE signal was at high levels in the vasculature and was visible within the LRP, with a brighter signal directly above the LRP (Figure 7A). We quantified fluorescence in the primordia with a region of interest drawn from the vasculature to the region directly above the primordia, and the fluorescence intensity was plotted as a function of the distance from the vasculature (Figure 7A). The signal directly above the LRP is significantly higher in the *tt7-2* mutant, mirroring the increased signal above the LRP observed using DCF, although *tt4-2* is not significantly different from Col-0. The signal throughout the LRP in *tt4-2* is comparable to wild type, while the signal in *tt7*-2 is significantly reduced, consistent with the presence of higher levels of kaempferol in *tt7-2* LRP. Therefore, we propose the decreased levels of superoxide detected by DHE within the primordia of *tt7-2* leads to impaired lateral root emergence.

**Figure 7:**
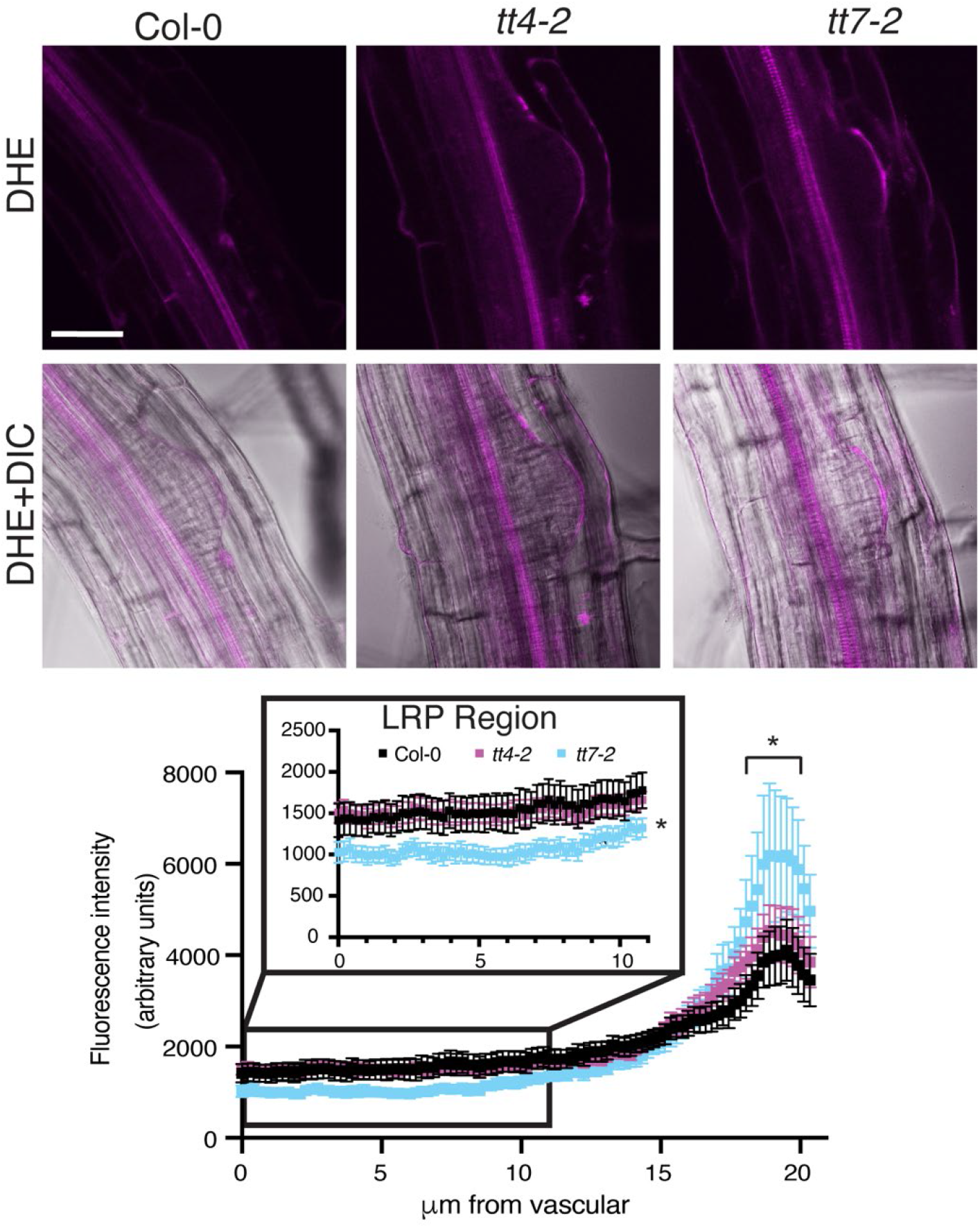
The tt7 mutant has decreased DHE signal within the lateral root primordia. 8-day old seedlings were stained with DHE and immediately imaged at the same using the same image settings over 3 separate experiments. DHE signal was quantified using FIJI and an 11.5μm was drawn from the base of the primordia to the tip of the primordia and the average and SE for 19-22 roots are reported. Statistics were determined using a one-way ANOVA based on the average of each line (*p<0.05).

### Treatment with IAA increases lateral root number for all flavonoid mutants

Since auxin drives initiation of LRP in pericycle cells and mutants with impaired flavonol synthesis have increased auxin transport (11, 17, 18), we tested an alternative hypothesis, that increased LRP formation in flavonol mutants was due to increased auxin delivery to pericycle cells. To examine whether the effect of flavonols on root development was through altered auxin transport, we transferred 5-day old seedlings of *tt4-2, tt4-11, tt4-11(CHS-GFP)*, and *tt7-2* mutants to media containing 0.1 μM or 0.3 μM indole-3-acetic acid (IAA) for three days. Comparison of the number of lateral roots revealed that all mutants and the wild type had increased number of emerged lateral roots after global treatment with IAA (Figure 8A), resulting in equivalent number of LRs in all genotypes. These data could be interpreted as the *tt4* mutants being less sensitive to IAA or having maximal auxin delivery to sites of lateral root initiation. Overall, this data is not consistent with differences in auxin transport or signaling controlling the differences in lateral roots between flavonol biosynthetic mutants.

**Figure 8:**
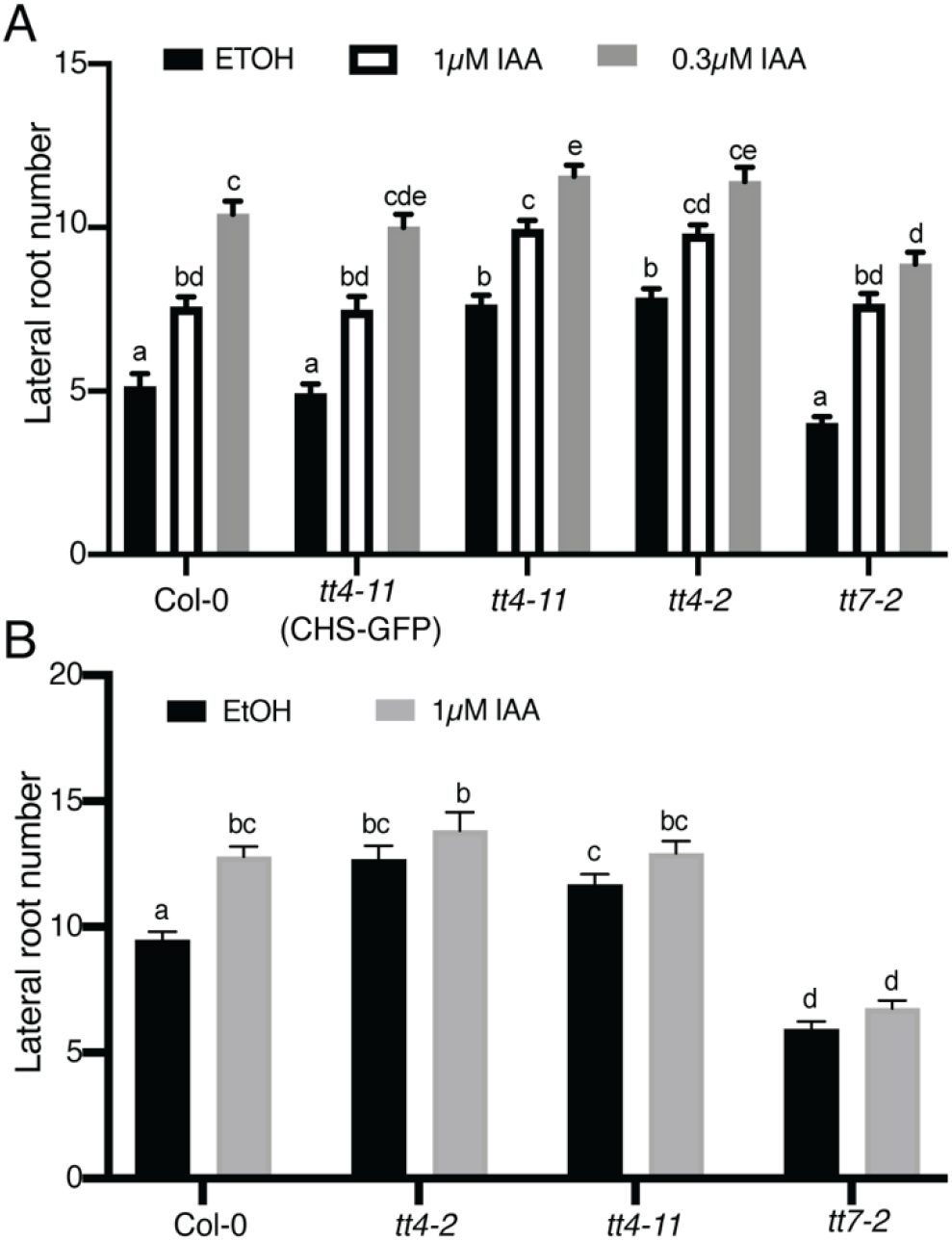
Flavonol mutants show maximal auxin delivery to the root. (A) Lateral root number was quantified for 8-day old seedlings after a 3-day IAA treatment across 3 individual experiments for a total n of: Col-0 (control: n= 30; 0.1μM IAA: n=30; 0.3μM IAA: n=30), *tt4-11*(CHS-GFP) (control: n= 39; 0.1μM IAA: n=40; 0.3μM IAA: n=39), *tt4-2* (control: n= 39; 0.1μM IAA: n=40; 0.3μM IAA: n=40), *tt4-11* (control: n= 40; 0.1μM IAA: n=40; 0.3μM IAA: n=40), and *tt7-2* (control: n= 40; 0.1μM IAA: n=40; 0.3μM IAA: n=40). (B) IAA containing agar droplets were placed at the root-shoot junction of 5-day old seedlings. Lateral root number was quantified for 9-day old seedlings across 6 individual experiments for a total n of: Col-0 (control: n= 69; IAA: n=75), *tt4-2* (control: n= 44; IAA: n=45), *tt4-11* (control: n= 69; IAA: n=70), and *tt7-2* (control: n= 60; IAA: n=65). Statistics were performed using a two-way ANOVA with a Tukey post hoc test. Bars with the same letter represent no statistical difference, while different letters indicate values that are significantly different with a p< 0.05.

To further test the effect of auxin on lateral root emergence in the flavonol mutants, we used a localized IAA treatment at the root shoot junction and measured the number of emerged lateral roots. Agar droplets containing either 1μM IAA in EtOH or EtOH were placed at the root-shoot junction of 5-day old seedlings and lateral root number was counted after 4 days of treatment. This treatment significantly increased lateral root number in Col-0 (Figure 8B), consistent with rootward transport of auxin stimulating lateral root emergence (61, 62). In contrast to wild type, the number of lateral roots was not significantly different in the presence or absence of IAA in *tt4-2, tt4-11*, or *tt7-2* (Figure 8B). These data are consistent with maximal auxin delivery in the absence of exogenous auxin in both *tt4* alleles and *tt7-2* and cannot account for opposite lateral root phenotypes between *tt4* and *tt7* alleles.

## Discussion

Flavonols are a subclass of flavonoid specialized metabolites that function as antioxidants in plants and regulate developmental and stress responses (2, 9, 15). Arabidopsis produces three flavonols, kaempferol, quercetin, and isorhamnetin (2, 63), which can be further modified via glycosylation, leading to a set of molecules with different antioxidant capabilities and distinct functions *in vivo* (5, 9, 13). The *tt4*(*2YY6*) and *tt4-1* mutants, which produce no flavonols, were previously reported to have increased numbers of lateral roots (18, 19). This study used a genetic approach to determine which flavonol is important for lateral root development and whether these biosynthetic and developmental defects were reversible using genetic and chemical complementation. Furthermore, we asked whether the alteration in root development in flavonol biosynthetic mutants is tied to their role as antioxidants.

Our experiments provide evidence that kaempferol is a negative modulator of lateral root development. We examined the lateral root number of a suite of flavonoid biosynthesis mutants and verified that each mutant had the predicted flavonol accumulation. In the absence of all flavonols (*tt4-2, tt4-11*, and *fls1-3*), there was an increase in lateral root number. This phenotype was reversed by genetic and chemical complementation. To eliminate the possibility that downstream anthocyanin metabolites control lateral root development, we used the *tt3* mutant that does not synthesize anthocyanins and found wild type levels of emerged lateral roots. Finally, we investigated which particular flavonol(s) is altering lateral root emergence using *omt-1*, which has reduced isorhamnetin concentrations and wild type kaempferol and quercetin, and *tt7-2* that is unable to produce quercetin or isorhamnetin and produces elevated levels of kaempferol. The *omt-1* mutant had subtle differences in total flavonols as there is limited conversion to isorhamnetin in Arabidopsis roots, and this mutant had wild type levels of emerged lateral roots. The most striking result was found with *tt7-2* which had elevated levels of kaempferol and significantly decreased numbers of emerged lateral roots. The lateral root phenotype of *tt7-2* suggests increased kaempferol concentration negatively impacts lateral root development, since the total flavonol level in this mutant is comparable to wild type.

We asked whether *tt4* and *tt7* mutations affected lateral root initiation and emergence. Altered lateral root number can be explained by either decreased emergence, reduced initiation, or a combination of the two. The *tt4* mutants had wild type levels of LRPs and increased levels of emerged lateral roots. In contrast, *tt7-2* formed increased numbers of LRPs, but had reduced numbers of emerged LRs. We also evaluated initiation using a root bending assay. There was significantly increased lateral root initiation in the *tt4* mutants, but the *tt7-2* mutant was comparable to wild type. These results are consistent with the elevated kaempferol in *tt7-2* inhibiting lateral root emergence, not initiation.

To understand why there is a reduced level of lateral roots in *tt7*-2, we examined the localization patterns of kaempferol and quercetin using the flavonol selective dye DPBA. Kaempferol accumulates in the LRP while quercetin has a more dispersed distribution in wild type roots. Within the LRP, kaempferol-DPBA fluorescence is increased in *tt7-2* compared to wild type where it is appropriately localized to impact lateral root development. The expression of a GFP reporter fusion to FLS suggests that flavonols are synthesized within lateral root primordia consistent with flavonol localization patterns determined using DPBA. The expression of the CHS-GFP fusion was limited in the primordia suggesting that there is transport of precursors into developing LRP. Consistent with this possibility, prior research has shown that naringenin can be transported long distances from the shoot to the root and vice versa, although flavonols do not move between tissues (16, 64). What is evident from these results is that there is precise spatial localization of flavonol accumulation, with different patterns found for quercetin and kaempferol. These findings suggest that the localization of the F3’H, which converts dihydrokaempferol to dihydroquercetin, may drive the localization of kaempferol to the LRP.

We tested the hypothesis that the enhanced lateral root formation in the absence of flavonols was tied to altered auxin transport, but our results do not support this hypothesis. The absence of flavonol metabolites leads to increased auxin transport in roots in both the shootward and rootward directions (11, 17, 18) and auxin drives lateral root formation (65, 66). We asked whether global or local treatments with IAA result in different responses in flavonol mutants. There was an induction of lateral roots in all mutants and wild type in response to global treatment with IAA, but no differences that suggest flavonols modulate the response to auxin. We also treated seedlings at the root-shoot junction to increase the available auxin for rootward transport and evaluated the lateral root number, since this polarity of auxin transport drives lateral root emergence (61, 62). This treatment increased the number of lateral roots formed in Col-0, but the *tt4* mutants and *tt7*-2 showed no significant change. These results are consistent with elevated auxin transport in *tt4-2, tt4-11*, and *tt7-2* (11). These results are therefore inconsistent with the lateral root phenotypes in the flavonoid biosynthetic mutants being the direct result of altered auxin transport.

Our results better support the hypothesis that the function of flavonol metabolites is to scavenge ROS that modulates lateral root development. Flavonol compounds have been shown to act as antioxidants *in vivo* to modulate stomatal aperture (10, 14), pollen viability (15), and root hair number (9). To ask if the developmental effects of flavonol mutants were tied to the ROS status, we treated roots with ascorbic acid. As an antioxidant, ascorbic acid has been shown to scavenge radical species (67) and is involved in reduction of ascorbate peroxidases that scavenge H_2_O_2_ (68). This antioxidant reduced the number of lateral roots in the *tt4* mutants to wild type levels but had no effect on root development in *tt7-2*, likely due to elevated levels of the flavonol antioxidant, kaempferol, in developing lateral roots. These results suggest that in *tt4* mutants elevated ROS drives the increase in lateral roots, while the increased kaempferol in LRP is sufficient to reduce ROS to levels that impair lateral root formation.

We worked to identify whether the ROS levels within the LRP or in tissues through which lateral roots emerge are altered in flavonol mutants to modulate LR emergence. We visualized patterns of ROS accumulation using a general ROS sensor DCF, and a superoxide sensor DHE. In wild type roots, DCF and DHE fluorescence are highest in roots above the LRP. The *tt4* mutants and *tt7-2* have an increased DCF and DHE fluorescent signal above the LRP compared to wild type. The elevated signal above the lateral root may be due to the absence of quercetin in this tissue in both mutants. Quercetin is a potent flavonol antioxidant, which in wild type is distributed throughout the root including in the layers overlying the primordia. The *tt4* mutants and *tt7-2* have the opposite lateral root phenotypes and similar ROS signals above primordia suggesting that the level of ROS in this region does not account for this developmental change. These results suggest the level of ROS above the LRP is not defining the differences in lateral root development in *tt4* and *tt7-2*.

The accumulation of superoxide within the LRP, as judged by DHE signal, is a more logical target of flavonol-modulated lateral root development. DHE fluorescence, but not DCF, is detected within the LRP. The different localization of these ROS sensors may be due to the reversible oxidation and signal of DCF leading to a transient signal (60) or to the selectivity of DHE for superoxide (69). Within the LRP, the DHE signal between Col-0 and *tt4-2* was equivalent but was significantly decreased in *tt7-2* compared to wild type. The reduction of DHE signal within the LRP of *tt7-2* is consistent with the 2-fold increases in kaempferol accumulation within primordia as detected by DPBA and across the entire root by LC-MS.

An important question is what leads to the localized superoxide accumulation within lateral root primordia and how does this ROS act to regulate development. Superoxide compounds can be produced by respiratory burst oxidase homologs (RBOHs) and then can be converted to hydrogen peroxide via superoxide dismutase (70). RBOHA-F localize in lateral root primordia determined using a GUS reporter (20). RBOHB and RBOHF localize at the base and the tip of the lateral root primordium, while the others localize throughout the primordium (20). The RBOH enzymes are therefore appropriately positioned to control the synthesis of superoxide in the developing lateral root primordia.

Another important question is how the superoxide in the LRP are able to modify lateral root development. This molecule may directly regulate the activity of proteins or mechanical properties of cell wall polymers or superoxide derived hydrogen peroxide may oxidize cysteine residues on proteins to alter their structure, activity, or function (71). A study identifying proteins that contain an oxidative modification in Arabidopsis found 4% of detected proteins were enriched in cell wall proteins (72). For instance, RBOHF has been shown to be regulate lignification of plant cell walls (73). Hydrogen peroxide has been implicated in strengthening of the cell wall through crosslinking of cell wall polymers (74). In contrast, hydroxyl radicals may loosen the cell wall (75). Cell wall structure is integral to lateral root emergence since the LRP must move through multiple cell layers requiring tightly controlled cell wall remodeling and mutants with impaired cell wall modifying enzymes have altered development (76–78). The cells within the primordia need to have strong cell wall structure to push through cell layers, and the cells overlying the primordia need to have their cell wall loosened to allow for emergence (76, 78). Therefore, one possible scenario is the decrease in the superoxide pool within the LRP in *tt7-2* may contribute to a less dense cell wall within the LRP impairing lateral root emergence.

The upregulation of flavonol biosynthesis is well described in response to environmental stress, hormones, and development signals (2, 9, 10, 14, 15). Additionally, flavonoid precursors can be transported both short and long distances (58, 59), which could implicate a role for flavonols in long distance signaling. Several recent studies have illustrated important roles of long-distance root to shoot signal transport in controlling root development (79). The HY5 transcription factor named for its control of hypocotyl elongation, also regulates primary root elongation and lateral root emergence, via mediating interactions between light and auxin signaling. Light-dependent synthesis of flavonols is positively regulated by this transcription factor (80). HY5 was suggested to move as a long-distance signal from the shoot to the root, but recent findings suggest HY5 controls synthesis of this mobile signal. An intriguing possibility is that HY5 regulated flavonoid precursors produced in the shoot tissue may act as a long-distance signal leading to altered root architecture due to shoot mediated environmental signals. We focused our study on signaling molecules downstream of flavonols, while signaling molecules upstream of flavonols that lead to root architectural changes are an interesting area of future study.

Lateral root development is a complex process that has multiple regulatory molecules. This study focuses on flavonol-modulated ROS within the lateral root primordia regulating lateral root emergence. We found that in the absence of flavonols, there is an increase in the number of lateral roots, which can be rescued with genetic and chemical complementation. Interestingly in the presence of increased kaempferol levels and no quercetin, as in *tt7-2*, there is a reduction of lateral root emergence. Kaempferol localizes primarily to vascular tissue and the lateral root primordia supporting its role in lateral root development. Flavonols have antioxidant properties which led us to evaluate the ROS concentration in and around LRP. We found the ROS concentration above the lateral root primordia was increased in the absence of quercetin (*tt4-2* and *tt7-2*) using DCF indicating it was not modulating flavonol-dependent lateral root development. We then used DHE to investigate the superoxide concentration in and around lateral root primordia. In *tt7-2* there was a reduction in DHE signal within LRP compared to wild type and *tt4-2*. Overall the data presented in this study suggests increased kaempferol within the lateral root primordia scavenge superoxide, leading to decreased DHE signal and impairing lateral root emergence.

## Experimental procedures

### Seeds and Growth Conditions

Arabidopsis mutants used in this study are in the Col-0 background. The following lines been previously published as follows: *tt4-2* (9, 81), *tt4-11* (11, 82), *fls1-3* (64), *tt7-2* (11), *CHS-GFP* (11), *FLS-GFP* (64), *omt-1* (63, 83). The *tt3* (SALK_099848C) mutant was obtained from the Arabidopsis Biological Resource Center. The T-DNA insert in the *DFR* gene in the *tt3* mutant was verified by PCR, shown to be the only insert, and homozygous lines were selected.

Arabidopsis thaliana seeds were sterilized in 90 % ethanol for five minutes and dried before sowing on 100 x 15 mm petri dishes with 30mL of 1X Murashige and Skoog (MS) medium (Caisson Labs), pH 5.5, with MS vitamins, 0.8% agar (MP Biomedicals), buffered with 0.05% MES (RPI), and supplemented with 1.5% sucrose. The petri dishes were sealed with micropore tape (3M). Seeds were stratified for 48hrs in darkness before being placed in vertical racks under continuous cool white light at 100-120 μmole photons m^-2^s^-1^. After five days, seedlings were transferred to either control or treatment plates and allowed to grow for three days. Plants treated with IAA and their controls were grown under a light bank with a yellow plexi-glass cover to reduce the amount of blue light exposure and reduce degradation of IAA (84). All assays took place three days after transfer.

### Lateral root imaging and quantification

Seedlings (8-day old) were imaged in brightfield mode using a Zeiss Axio Zoom V16 stereomicroscope with a Zeiss AxioCam 506 monochrome camera using panorama mode. Lateral root numbers were quantified in 8-day old seedlings using an Olympus SZ40 microscope. Statistical analysis of the data was completed in Prism 8 using a two-way ANOVA followed by Tukey’s post-hoc test.

For treatments with naringenin or ascorbic acid, seedlings were 5-days old at the time of treatment and lateral root number was quantified three days after treatment. MS media was prepared as described above and was cooled for 1 hour in a 60°C water bath after autoclaving before treatments were added to the media. Stock solutions of treatments were prepared for each individual experiment. Naringenin (Indofine Chemicals) was prepared as a 250 mM stock in EtOH and was diluted in the media to a final concentration of 100 μM. Ascorbic acid (Sigma) was prepared as a 1M stock in water and diluted in the media to a final concentration of 500 μM.

Seedlings treated with IAA were five days old at the time of treatment and lateral root number was quantified after three days of treatment. MS media was prepared as described previously and cooled for 1hr at 55°C in a water bath after autoclaving before treatments were added to the media. A stock solution of 10mM IAA in EtOH was diluted to 100μM in EtOH and then diluted further within the media to final concentrations of 0.1μM and 0.3μM. Treatment with auxin at the root-shoot junction occurred on 5-day old seedlings transferred to MS media. Agar droplets were made using 0.8% agar and allowed to cool for 20 minutes. A stock solution of 10mM IAA was diluted to 100μM and then diluted further within the agar to final concentration of 1μM. A pipette was used to measure out 30μl of the agar with IAA or EtOH and expelled in a droplet on a strip of parafilm. These droplets cooled and solidified for 20 minutes, before being placed at the root-shoot junction of 5-day old seedlings. Lateral root number was quantified 4 days after treatment.

### Clearing and quantification of lateral root primordia

To quantify the number of LRP, seedlings were fixed and cleared with the Clearsee method (85). Roots were fixed in 4% paraformaldehyde (Acros organics) in PBS (Alfa Aesar) for 1 day. Seedlings were rinsed twice for one minute in PBS buffer before being transferred to the Clearsee solution (10% xylitol, 15% sodium deoxycholate, 25% urea in water). Seedlings were incubated in Clearsee for 2-3 days prior to being mounted on microscope slides in water. The number of lateral root primordia was counted using brightfield mode of a Zeiss Axio Zoom V16 stereomicroscope with a Zeiss AxioCam 506 monochrome camera. The number of lateral roots per seedling are reported. Statistical analysis of the data was completed in Prism 8 using a two-way ANOVA followed by a Tukey’s post-hoc test.

LRP were also quantified using a root bending assay. Seedlings were grown to five days old and transferred to control plates. Each seedling was bent 1-2mm at the tip to a 90-degree angle of bending at the root at X distance from the tip. The presence of a LRP or an emerged lateral root was quantified two days after bending (48hrs). Emerged lateral roots were quantified using an Olympus SZ40 microscope. Seedlings were then imaged under a REVOLVE microscope with a drop of water placed on the bend area to determine if there was a primordium. The formation of a lateral root (emerged or unemerged) is reported in the form of a ratio to the number of roots bent. Statistical analysis of the data was completed in Prism 8 using an ordinary one-way ANOVA followed by a Dunnett’s multiple comparisons test.

### DPBA imaging and quantification

Diphenylboric acid (DPBA, Sigma Aldrich) was used to visualize quercetin and kaempferol *in vivo* using previously published methods that use slight signal differences in emission spectra between the two flavonols to elucidate their unique accumulation patterns (11). Seedlings were stained in darkness in 0.25% w/v DPBA with 0.06% Triton-X (v/v) in water for 12 minutes, with a 12 minutes wash in water. Seedlings were mounted on microscope slides in water. Roots were imaged using a Zeiss LSM 880 confocal microscope using the 458 laser at 8% laser power. Images were taken using a 40X objective with an emission spectrum 472-499 for the kaempferol channel and 585-619 for the quercetin channel. Z-stack images were taken to identify the center of the lateral root primordia. All microscope settings were identical for each biological replicate.

DPBA fluorescence was quantified for each lateral root primordia using FIJI. The center of the lateral root primordia was determined, and the channels were pseudo-colored green for kaempferol and yellow for quercetin. A 100-pixel line (which was 11.5 μm) was then drawn from the vascular tissue to the epidermis through the center of the primordia. The fluorescence intensity was measured on average every 0.1μm along the line and measurements were taken from 0 to 60 μm across the lateral root primordia. A single optical slice was used for quantification and as a representative image.

### *CHSpro:CHS:GFP* and *FLSpro:FLS:GFP* confocal imaging

8-day old seedlings transformed with the *CHSpro:CHS:GFP* (11) or the *FLSpro:FLS:GFP* (64) reporters were imaged using the Zeiss LSM 880 confocal microscope and the 488nm laser at 12% power and 490-544 emission spectra. Cell walls were stained with 0.5mg mL^-1^ propidium iodide (PI, Acros Organics) dissolved in water for 5 minutes. PI was imaged by excitation with a 561 laser with an emission spectrum set to 579-695nm. All microscope settings were identical for each biological replicate. Z-stacks were taken and compressed into maximum intensity projections using Zen black software.

### DCF and DHE imaging and quantification

General ROS were detected using 2’-7’-dichlorodihydrofluorescein diacetate (CM-H_2_DCFDA, Invitrogen). CM-H_2_DCFDA was dissolved in DMSO to give a concentration of 1 mM and further diluted to a final concentration of 50 μM in water. Both control and ascorbic acid treated seedlings were incubated in CM-H-2DCFDA for 12 minutes and washed in water for 1 minute in the dark before being mounted on microscope slides in water. Fluorescence was detected using a Zeiss LSM 880 confocal microscope using the 488nm laser at 5% power with and an emission spectrum of 490-606nm. For each lateral root primordia, z-stack images were generated using the 40X objective with a line averaging of 4. We were careful to minimize photooxidation by using a lower laser power and limiting exposure of the sample to the laser and light. Similarly, we obtained images swiftly after exposure to DCF to limit any photooxidation and changes to the ROS environment that were not related to our experiment.

To verify that the dye was being taken up into the primordia, fluorescein diacetate (FDA, Acros organics) controls were performed. FDA was prepared and imaged in the same manner as CM-H_2_DCFDA. FDA was not taken into primordia of stage 3 and lower, therefore those primordia were not included in the quantification. Zen Blue software was used for quantification of DCF fluorescence. A region of interest was drawn along the epidermis above and below the lateral root primordia. The region of interest varied slightly based on size of the lateral root primordia but averaged to 3,185.7μm^2^ for above the primordia and 3,010.1μm^2^ below the primordia. Statistical analysis of the data was completed in Prism 8 using a two-way ANOVA followed by Tukey’s post-hoc test.

Superoxide accumulation was visualized using dihydroethidium (DHE). DHE was dissolved in DMSO to a concentration of 20 mM and further diluted to 50 μM in water. 8-day old seedlings were incubated in DHE for 25 minutes and then rinsed in water before being mounted on microscope slides. Fluorescence was detected using a Zeiss 880 confocal microscope using the 488 laser at 2% power with an emission spectrum of 490 to 561 nm. For each lateral root primordia, Z-stacks were generated using the 40x water objective with a line averaging of 4. FIJI was used for image quantification. A 100-pixel line (11.5μm) was drawn from the vascular tissue to the tip of the lateral root primordia. Statistical analysis was completed in prism 8 using a two-way ANOVA followed by Tukey’s post-hoc test.

### Extraction and quantification of aglycone flavonoids by LCMS

Flavonols were extracted as previously described (9). Roots of 7-day old seedlings were separated from shoots and flash frozen in liquid nitrogen and immediately used for extraction or stored at −80°C until extraction. The extraction buffer had an internal standard of 500nM formononetin (Indofine chemicals) that was dissolved in 100% acetone (Optima-grade, Fisher Scientific). The extraction buffer was added to the samples, which ranged between 17.65mg and 65.01 mg, at 3μL/mg. Tissues were homogenized using a 1600 MiniG tissue homogenizer (Spex Sample Prep) for 10 minutes. An equal volume of 2N HCl was added to the samples and they were incubated at 75° C for 45 minutes to produce aglycone flavonols. 300 μL of ethyl acetate (Optima grade, Fisher) was added to the samples and shaken for five minutes before centrifugation at maximum speed for 10 minutes. The top organic phase was isolated and the ethyl acetate phase separation was repeated and the organic phases were pooled. The organic phase was dried by airflow using a mini-vap evaporator (Supelco Inc). Samples were resuspended in 300μL of acetone before LCMS analysis.

Samples were analyzed on a Thermo LTQ Orbitrap XL with an electrospray ionization source, coupled to a Thermo Accela 1250 pump and autosampler (Thermo Fisher). For flavonols analysis, 10μL of each samples was injected with a solvent of water: acetonitrile, both containing 0.1% (v/v) formic acid on a Luna 150 x 3mm C18 column with a Security guard pre column. The solvent gradient was 90% water: 10% acetonitrile (v/v) to 10% water: 90% acetonitrile (v/v) in a time span of 18.5 minutes. From 18.5 minutes to 20 minutes, the gradient moves from 10% water: 90% acetonitrile (v/v) back to 90% water: 10% acetonitrile (v/v) and holds at these concentrations for another 2 minutes to recondition the column. Quantification of flavonols was found by comparing peak area data, quantified in Thermo Xcalibur, to standard curves generated using pure standards of naringenin, quercetin, kaempferol, and isorhamnetin (Indofine chemicals). Total flavonol level was calculated by averaging the total flavonol value of each sample. MS2 fragmentation spectra were induced using 35-kV collision-induced dissociation and the MS2 spectra of flavonols was compared to standards in Massbank database. Statistical analysis of the data was completed in Prism 8 using a two-way ANOVA followed by a Tukey’s post-hoc test.

## Data availability

All data are included in this article.

## Acknowledgements

We appreciate the assistance of Heather Brown Harding (Microscopy core facility) with imaging and Christopher Tracy (Mass spectroscopy facility) with development of LC-MS methods and analysis approaches. The helpful editorial suggestions of Joelle Muhlemann are also appreciated.

## Abbreviations

ROS: Reactive oxygen species
CHS: Chalcone synthase
LRP: Lateral root primordium
LC-MS: Liquid chromatography-mass spectrometry
FLS1: Flavonol synthase 1
F3’H: Flavonoid 3’hydroxylase
DFR: Dihydroflavonol 4-reductase
DPBA: Diphenylboric acid 2-aminoethyl ester
LSCM: Laser scanning confocal microscopy
DIC: Differential interference contrast
CM H_2_DCF-DA: 2’,7’-dichlorodihydro-fluorescein diacetate
FDA: Fluorescein diacetate
DHE: Dihydroethidium
IAA: Indole-3-acetic acid
RBOH: Respiratory burst oxidase homologs
CHI: Chalcone isomerase
F3H: Flavonoid 3 hydroxylase
OMT-1: O-methyl transferase 1

## Supplemental Figures

**S1:**
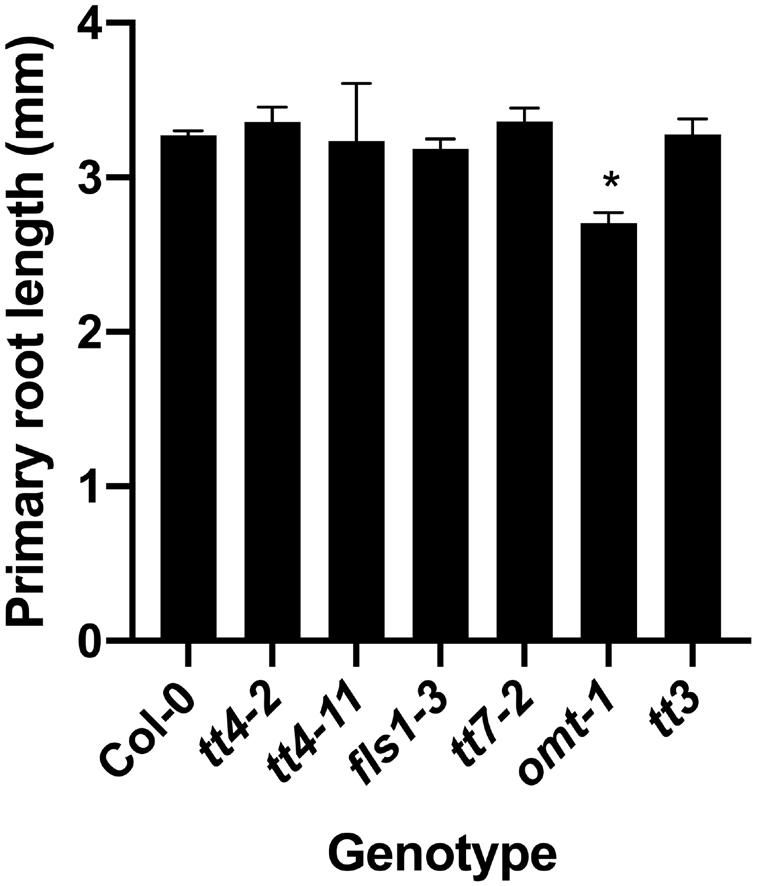
Primary root length is similar in all mutants except *omt-1*, which has significantly shorter primary roots. Primary root length was measured in 8-day old seedlings across X individual replicates for a total n of X. Statistics were measured using a one-way ANOVA followed by a Dunnett’s multiple comparisons test. (*p<0.0001).

**S2:**
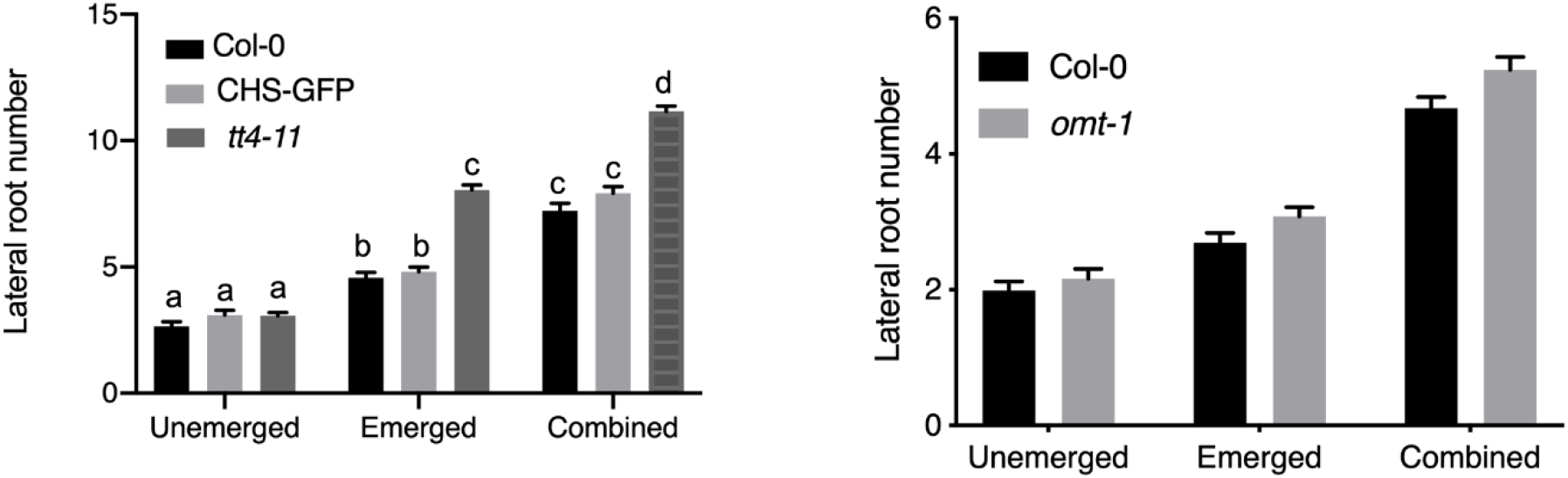
Lateral root primordia are comparable between wild type and *omt-1*. The number of lateral primordia (Stages 4-7) and emerged lateral roots, and the combined totals were quantified in cleared 7-day old seedlings. The average and SE from four separate experiments with a total n=60-69. Statistical differences were determined with a two-way ANOVA with a Tukey’s post hoc test. Bars with the same letter represent no statistical difference, while different letters indicate values that are significantly different with a p< 0.05.

**S3:**
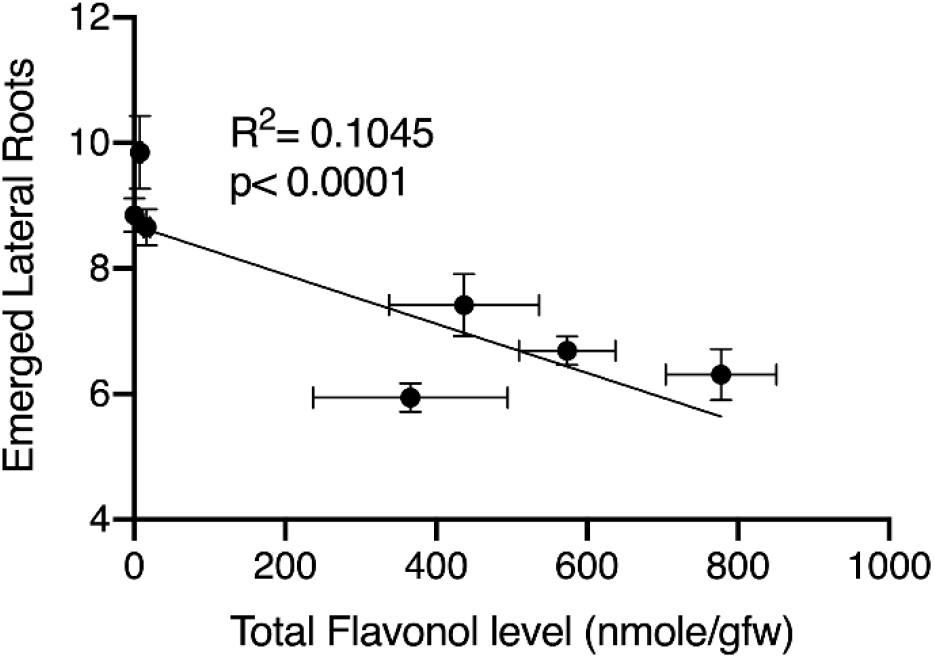
Correlation between total flavonol levels and emerged lateral roots for flavonoid biosynthesis mutants. The number of emerged lateral roots was plotted as a function of the total flavonol level determined by LCMS for each flavonoid biosynthesis mutant. A simple linear regression was performed and the correlation and p-values reported.

**S4:**
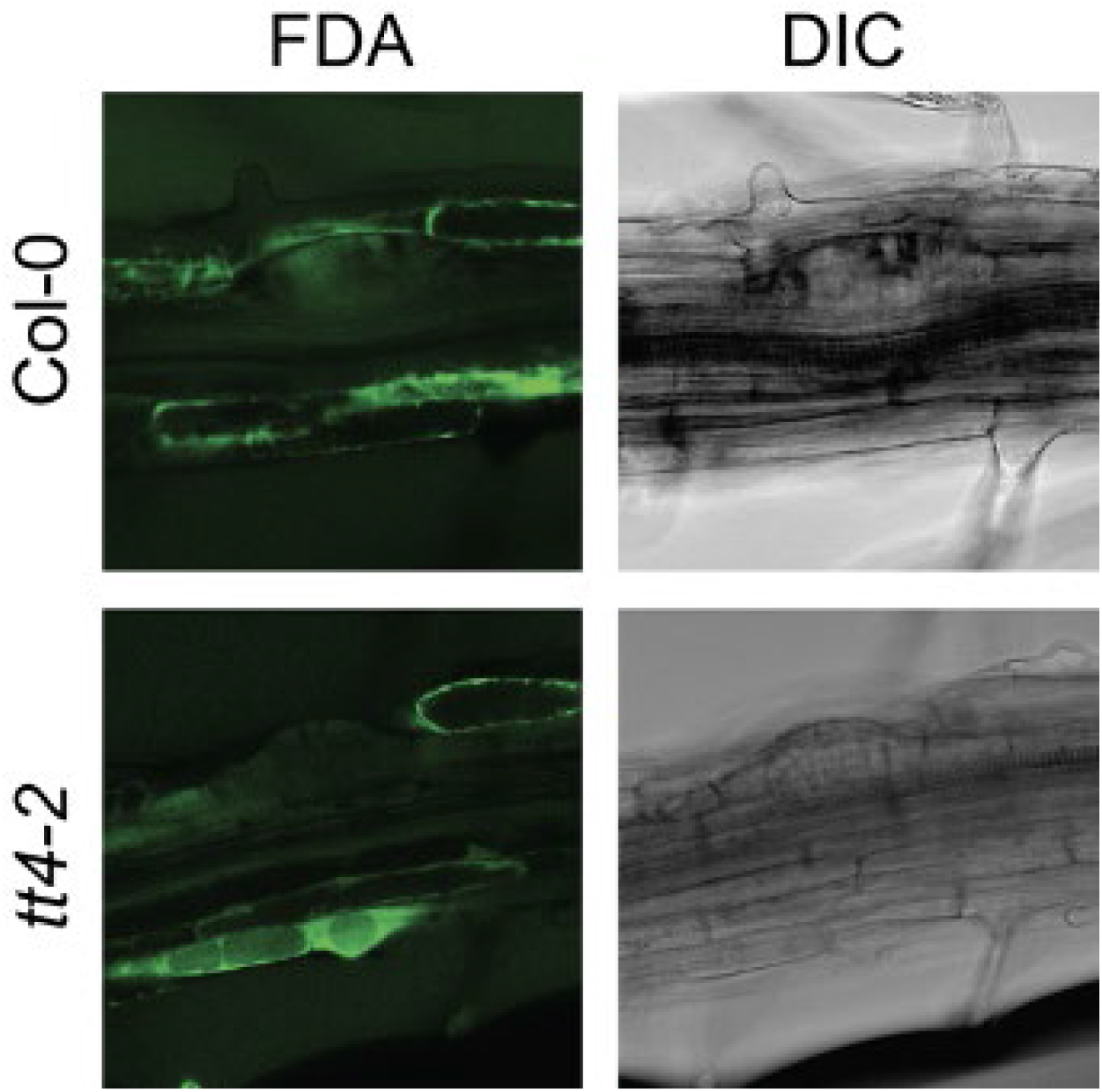
FDA is able to enter later stage lateral root primordia. 8-day old seedlings were stained with FDA and imaged using the same settings across 2 replicates. LRP from stage 2 to stage 8 were evaluated based on their ability to take up FDA. LRP stage 3 and below were unable to take up the FDA. Therefore, experiments using DCF only evaluated stage 4 and higher LRP.

**S5:**
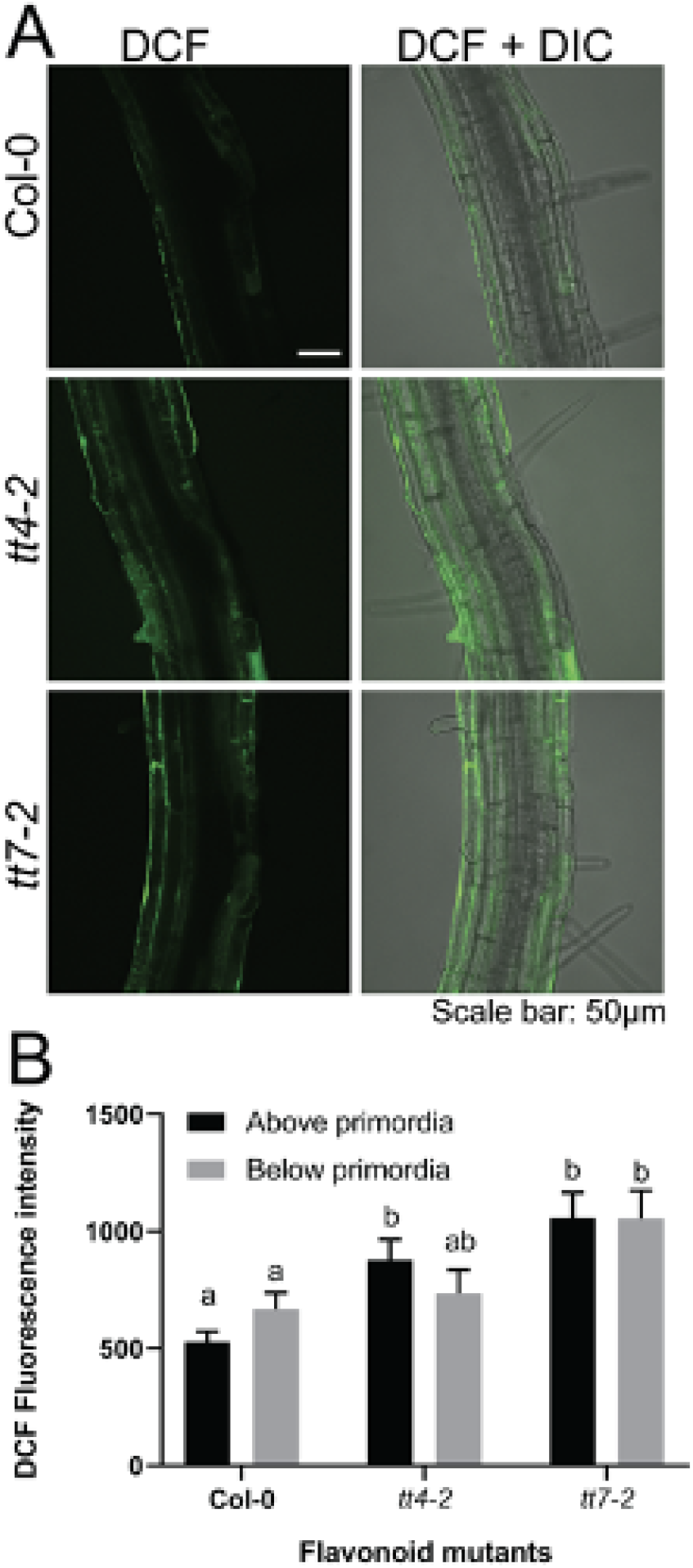
Mutants with reduced flavonoid biosynthesis have increased DCF signal. 8-day old seedlings were stained with DCF and immediately imaged at the same laser power, gain, and pinhole for six separate experiments. DCF was imaged in stage 4 and lateral roots. DCF signal was quantified using Zen blue edition and a region of interest in the epidermis above and below lateral root primordia. Statistics were determined using a two-way ANOVA with a Tukey post-hoc test for a total n of Col-0 (n=39), *tt4-2* (n=34), *tt7-2* (n=24). Bars with the same letter indicate no statistical significance while bars with different letters are statistically significant from one another. P<0.05

